# An amplicon panel for high-throughput and low-cost genotyping of Pacific oyster

**DOI:** 10.1101/2023.08.26.554329

**Authors:** Ben J. G. Sutherland, Neil F. Thompson, Liam B. Surry, Krishna Reddy Gujjula, Claudio D. Carrasco, Srinivas Chadaram, Spencer L. Lunda, Christopher J. Langdon, Amy M. Chan, Curtis A. Suttle, Timothy J. Green

**Affiliations:** Sutherland Bioinformatics, Lantzville, BC, Canada V0R 2H0; Faculty of Science and Technology, Vancouver Island University, Nanaimo, BC, Canada V9R 5S5; Pacific Shellfish Research Unit, Agricultural Research Service, United States Department of Agriculture, Hatfield Marine Science Center, Newport, OR 97365 USA; ThermoFisher Scientific, 2130 Woodward Street, Austin, TX 78744 USA; Oregon State University, Department of Microbiology, 226 Nash Hall, Corvallis, OR 97331; Oregon State University, Coastal Oregon Marine Experiment Station, Hatfield Marine Science Center, 2030 SE Marine Science Dr., Newport, OR 97365; Department of Earth, Ocean and Atmospheric Sciences, The University of British Columbia, Vancouver, BC, Canada V6T 1Z4; Department of Microbiology and Immunology, The University of British Columbia, Vancouver, BC, Canada V6T 1Z3; Department of Botany, The University of British Columbia, Vancouver, BC, Canada V6T 1Z4; Institute for the Oceans and Fisheries, The University of British Columbia, Vancouver, BC, Canada, V6T 1Z4

**Keywords:** amplicon panel, aquaculture, *Crassostrea* (*Magallana*) *gigas*, genetic diversity, Pacific oyster, parentage assignment

## Abstract

Maintaining genetic diversity in cultured shellfish can be challenging due to high variance in individual reproductive success, founder effects, and rapid genetic drift, but is important to retain adaptive potential and avoid inbreeding depression. To support broodstock management and selective breeding in cultured Pacific oysters (*Crassostrea* (*Magallana*) *gigas*), we developed an amplicon panel targeting 592 genomic regions and SNP variants with an average of 50 amplicons per chromosome. Target SNPs were selected based on elevated observed heterozygosity or differentiation in Pacific oyster populations in British Columbia (BC), Canada. The use of the panel for parentage applications was evaluated by genotyping three generations of oysters from a breeding program in BC (n = 181) and a set of families that were selected for Ostreid herpesvirus-1 (OSHV-1) resistance from the Molluscan Broodstock Program, a Pacific oyster breeding program in Oregon, USA (n = 136). Population characterization was evaluated using collections of wild, naturalized, farmed, or hatchery oysters sampled throughout the Northern Hemisphere (n = 190). Technical replicate samples showed high genotype concordance (97.5%; n = 68 replicates). Initial parentage analysis found instances of suspected pedigree and sample handling errors, demonstrating the panel’s value for quality control in breeding programs. Suspected null alleles were identified in parentage datasets and were found to reduce assignment success. Null alleles were largely population dependent, suggesting population-specific variation impacts target amplification. By taking an iterative approach, null alleles were identified using existing data without the need for pedigree information, and once null alleles were removed, assignments increased to 93.0% and 86.0% of possible assignments in the two breeding program datasets. A pipeline for analyzing the amplicon sequence data from sequencer output, *amplitools*, is also provided.

## Introduction

Sustainable aquaculture depends on the effective characterization, preservation, exchange and use of genetic resources (Guo, 2009). High resolution genomic tools (e.g., high-density genetic maps, SNP chips, chromosome-level reference genomes) enable genome-wide association studies to identify markers that can be used for marker-assisted or genomic selection (Boudry et al., 2021). Low resolution genetic tools (e.g., amplicon panels) enable high-throughput genotyping at low cost (Meek & Larson, 2019). Lower-cost applications can allow scalable use, such as in fisheries management through parentage-based tagging (e.g., Beacham et al., 2018), or in aquaculture to assign individuals back to putative family when mixed without physical tagging (Allen et al., 2020). Amplicon panels can provide rapid turnaround time (Arbelaez et al., 2019), and increased flexibility to use lower quality input DNA (Csernák et al., 2017); in general, these panels have been described as reliable, scalable, moderately easy to analyze, and cost-effective (Meek & Larson, 2019). Cost-effectiveness can bring genetic advances to small or medium sized operations not able to afford high-density genotyping tools (Boudry et al., 2021; Delomas et al., 2023).

Pacific oyster *Crassostrea* (*Magallana*) *gigas* (Thunberg, 1793; Salvi & Mariottini, 2017), is one of the most valuable aquaculture species globally (Botta et al., 2020). Oyster aquaculture is a significant economic driver and food source for many developed and developing nations (Martínez-García et al., 2022). However, production and growth of the industry outside of China has stagnated due to disease, regulatory issues, and other factors (Botta et al., 2020). Large-scale mortality events have been a significant burden on the industry, where disease may be caused by specific or combined protozoan, bacterial, or viral pathogens, as well as by multifactorial or unexplained causes (King et al., 2019). Abiotic stressors such as salinity shifts, elevated temperatures, or heat waves can interact with these biotic stressors, increasing negative impacts (Green et al., 2019; King et al., 2019).

Environmental changes more generally are also challenging cultured and wild oysters. Ocean acidification negatively impacts shell formation and maintenance (Barros et al., 2013; Barton et al., 2012; Watson et al., 2009), and extreme weather has led to massive mortality events in recent years. For example, an atmospheric river and freshwater influx into the San Francisco Bay (California, USA) coincided with a near 100% mortality event of the highly abundant population of wild Olympia oyster *Ostrea lurida* (Cheng et al., 2016). An unprecedented extreme heatwave in the Pacific Northwest during summer 2021 resulted in large-scale negative impacts to wild and naturalized shellfish in the area (Raymond et al., 2022). With a multitude of stressors and impediments to industry stability and growth, shellfish farmers in the Pacific Northwest consider oyster health and selective breeding to be important scientific adaptation strategies to increase crop resilience (Green et al., 2023; Thompson, 2023). As such, improvements to selective breeding infrastructure and genomic resources are valuable to Pacific oyster breeders, as recently advocated for eastern oyster *C. virginica* (Allen et al., 2020).

Maintenance of genetic diversity is essential to shellfish breeding. Although standing polymorphism levels are very high in shellfish (Plough, 2016), where a SNP is expected every 40 bp in *C. gigas* (Hedgecock et al., 2005; Sauvage et al., 2007), rapid reductions in genetic diversity can occur in cultured lineages (Evans et al., 2004; Gurney-Smith et al., 2017; Hedgecock & Sly, 1990; Xiao et al., 2011), with losses compounding over time, leading to significant concerns for breeders (e.g., Li et al., 2007). High genetic load occurs through high fecundity and likely high mutation rates, which can produce severe inbreeding depression affecting growth and survival (Evans et al., 2004). Although inbreeding depression may be counteracted by crossing divergent lines to achieve heterosis (Hedgecock & Davis, 2007; Hedgecock et al., 1995), long-term retention of genetic diversity, and maintenance of hatchery lineages is important for broodstock programs (Carlsson et al., 2006; Gurney-Smith et al., 2017). Furthermore, retaining standing genetic diversity is vital for rapid adaptation (Barrett & Schluter, 2008) and selective breeding (Guo, 2009).

A rapid, inexpensive, automatable, and easily analyzed genotyping tool is needed for the Pacific oyster. Existing genomic resources for the Pacific oyster include high quality reference genome assemblies (Peñaloza et al., 2021; Qi et al., 2021) and genetic maps (Gutierrez et al., 2018; Li et al., 2018; Yin et al., 2020), as well as SNP arrays of high (134K targets; Qi et al., 2017), medium (41K; Gutierrez et al., 2017), and low density (384; Lapègue et al., 2014). However, due to significant costs and required expertise, smaller or medium-scale breeding operations may not be able to benefit from these resources (Boudry et al., 2021). Until present, there has been no low-density amplicon panel for the Pacific oyster. In addition to lower cost per individual, amplicon panels can also be used to genotype non-target variants within an amplicon window, potentially expanding its utility across populations and time, as long as primer sites remain intact and functional. This longevity and malleability may be particularly advantageous to Pacific oysters given their high genetic diversity (Hedgecock et al., 2005; Sauvage et al., 2007), global scale of culture (Boudry et al., 2021; Martínez-García et al., 2022), including genetically diverse large-scale commercial hatcheries (Sutherland et al., 2020), and temporal fluctuations in allele frequencies that occur due to sweepstakes reproductive success (Hedgecock, 1994; Hedgecock & Pudovkin, 2011; Sun & Hedgecock, 2017).

Here we describe the development of a 592 target SNP panel designed using previously identified high heterozygosity and high differentiation markers from naturalized Pacific oysters in British Columbia (BC), Canada, which are similar to other global populations derived from the Japan translocation lineage (Sutherland et al., 2020). We test the panel on samples from Canada, France, Japan, and China, as well as cultured populations from the United Kingdom, the United States, and Canada. We further evaluate and demonstrated the panel’s effectiveness for genetic parentage assignment using single SNP targets through analyzing three generations of the Vancouver Island University (VIU) breeding program, and a closely related set of families bred for viral resistance from Oregon State University’s Molluscan Broodstock Program (MBP). All design amplicons and target variant identities and sites are provided here, and the panel is available via the commercial provider (ThermoFisher Scientific). All analytic code is available in *amplitools*, a repository designed to move from sequencer output to parentage analysis (see *Data Availability*). Details on the latest version of the panel’s target file is available in the accompanying repository *amplitargets* (see *Data Availability*), expected to be updated over time. Collectively, with these outputs we aim to facilitate uptake and therefore bring rapid gains in Pacific oyster breeding and aquaculture.

## Methods

### Data sources and marker selection

Plink and Variant Call Format (VCF) outputs from Sutherland et al. (2020), a restriction-site associated DNA sequencing (RAD-seq) study of Pacific oysters from the Northern Hemisphere, were obtained from the authors and uploaded to FigShare (see *Data Availability*). Genotypes were read into R (R Core Team, 2023) using the read.PLINK function of adegenet (v.2.1.5; Jombart & Ahmed, 2011) and data were converted from genlight format to genind, genepop, and hierfstat formats for analysis using adegenet and hierfstat (v.0.5-10; Goudet & Jombart, 2022).

The pre-filtered dataset was limited to only include naturalized collections from British Columbia (BC), the focal region of the study, which includes the locations Hisnit (HIS), Pendrell (PEN), Pipestem (PIP), and Serpentine (SER); for details on collection sites see Sutherland et al. (2020). Within the BC dataset, per locus global minor allele frequency (MAF) was recalculated, and SNP variants with MAF < 0.01 were removed. Per locus observed heterozygosity (*H*_OBS_) and average *F*_ST_ (Weir & Cockerham, 1984) were calculated using adegenet and pegas (v.0.12; Paradis, 2010), respectively. SNPs were excluded from the RAD-seq dataset when *H*_OBS_ > 0.5 in an effort to avoid duplicated genomic regions, and remaining SNPs with the highest *H*_OBS_ (n = 300), or the highest *F*_ST_ (n = 300) were selected for panel design. Several additional SNP variants of interest were included based on previous detection as private alleles in Deep Bay, BC (n = 15) or Guernsey, UK (n = 5) farmed populations (Sutherland et al., 2020). Nine SNPs were redundant between the *F*_ST_ and *H*_OBS_ selected lists; 611 unique SNPs were identified for panel design (Table S1).

### Panel design and chromosomal locations

Flanking 200 bp sequences on either side of selected SNPs were obtained from the contig-level reference genome (GCA_000297895.1; Zhang et al., 2012) that was used in marker discovery (Sutherland et al., 2020) using a bed file derived from the marker discovery VCF and the bedtools function getfasta (Quinlan & Hall, 2010). All required code and instructions for this process are provided (see *Data Availability*, *ms_oyster_panel*). Target amplicon window sequences (401 bp), variant positions, and reference and alternate alleles (Additional File S1) were submitted to the AgriSeq primer design team for the Ion Torrent system (ThermoFisher Scientific). Targets were passed through the AgriSeq quality control process (ThermoFisher Scientific), using the GCA_000297895.1 genome. Those passing QC were then submitted to the primer design phase. In the design phase, oligo candidates were generated, scored, filtered, and the optimal oligo pair for each target was selected to be included in the final panel. The selected oligos were checked *in silico* for specificity and sensitivity of the intended target regions using the *C. gigas* reference genome GCA_000297895.1. The resultant amplicon panel is available commercially through ThermoFisher Scientific (SKU A58237 AGRISEQ PACIFIC OYSTER PANEL).

As the marker discovery RAD-seq genotyping (Sutherland et al., 2020) used a contig-level genome assembly (Zhang et al., 2012), the chromosomal positions of the targeted 401 bp sequences were subsequently determined by mapping them against a chromosome assembly (GCF_902806645.1; Peñaloza et al., 2021) using bowtie2 (Langmead & Salzberg, 2012) in end-to-end mode allowing multiple alignments (-k 6). The target sequences were also aligned with bwa mem (Li, 2013). Amplicons were categorized as mapping to a single or multiple positions, and those mapping to multiple positions were removed during analysis. Positions of amplicon targets along chromosomes were visualized using ggplot2 (Wickham, 2016).

### Panel testing

The panel was tested for use in characterizing different populations by genotyping oyster collections from 2017 and 2018, as described by Sutherland et al. (2020) and shown in Table 1, including from a farm (Deep Bay, BC, DPB), commercial hatcheries (Guernsey, UK, GUR; and China, QDC), and wild (Japan, JPN; China, CHN) or naturalized sources (Pendrell Sound, BC, PEN; France, FRA). Although these same populations and collection years were used for marker discovery (Sutherland et al., 2020), the panel testing was done on alternate individuals not used for discovery, except for JPN and CHN since additional samples were not available. Notably, JPN and CHN were not used for marker selection, as described above.

**Table 1.** The sources and types of collections used in the study are shown alongside the number of unique oysters genotyped per collection, and the number and percentage of samples retained after filtering. OFR1 and OFR2 are the broodstock for OFR5, and G0 are the broodstock for OFR1 and OFR2. Acronyms: VIU = Vancouver Island University; MBP CHR8 = Molluscan Broodstock Program chromosome 8 families; BC = British Columbia.

| Collection | Type | Country of origin | Country collected | Genotyped (n) | Filtered (n) | Retained (%) |
| --- | --- | --- | --- | --- | --- | --- |
| Pendrell (PEN) | Naturalized | Canada | Canada | 31 | 29 | 93.5% |
| France (FRA) | Naturalized | France | France | 32 | 31 | 96.9% |
| Japan (JPN) | Wild | Japan | Japan | 22 | 22 | 100.0% |
| China (CHN) | Wild | China | China | 33 | 33 | 100.0% |
| China (QDC) | Commercial hatchery | China | China | 8 | 8 | 100.0% |
| Guernsey (GUR) | Farm / commercial hatchery | United Kingdom | BC, Canada | 31 | 15 | 46.9% |
| Deep Bay (DPB) | Farm / commercial hatchery | *unknown* | BC, Canada | 32 | 32 | 100% |
| VIU G0 (F0) | University hatchery | *unknown* | BC, Canada | 16 | 11 | 68.8% |
| VIU OFR5 parents (F1) | University hatchery | *unknown* | BC, Canada | 53 | 40 | 75.5% |
| VIU OFR5 offspring (F2) | University hatchery | *unknown* | BC, Canada | 112 | 91 | 81.3% |
| MBP CHR8 F0 | University hatchery | *unknown* | USA | 25 | 24 | 96% |
| MBP CHR8 F1 | University hatchery | *unknown* | USA | 111 | 111 | 100% |
| **Totals** |  |  |  | 506 | 447 | 88.3% |

The panel was tested for utility in parentage analysis by genotyping samples from three generations of the Vancouver Island University (VIU) breeding program, and oyster families from two MBP generations that were selected based on the presence or absence of a SNP marker on chromosome 8 (CHR8) for resistance to the Ostreid herpesvirus-1 (OsHV-1) (Divilov et al., 2023). The breeding program samples were provided with information about their respective generation, but pedigree information was not used until after assignments were created and the initial analysis reviewed. These analyses are described below.

DNA from the VIU and MBP breeding program samples was extracted using the Monarch Genomic DNA Purification kit (NEB) using the enzymatic cleanup tissue extraction protocol with proteinase K and RNase A and quantified using a BioSpectrometer (Eppendorf). All other samples were extracted using the BioSprint (QIAGEN) method and diluted to 25 ng/μl and stored at −20°C. Samples were further normalized to 10 ng/μl in 25 μl volumes and submitted to ThermoFisher Scientific laboratories in Austin, TX for AmpliSeq library preparation using the AgriSeq™ HTS Library kit (ThermoFisher Scientific) with 16 cycles of amplification and IonCode barcode labelling. Libraries were sequenced using Ion 540 chips on an Ion Torrent S5 (ThermoFisher Scientific). Some samples had low sequence yields, and so a portion of the poorly amplifying libraries were synthesized again from undiluted source samples, then sequenced at both diluted and supplied sample concentrations. Replicate samples with sufficient genotyping rates per sample were used to evaluate repeatability of library preparation, sequencing, and genotyping (more details provided below). The MBP samples were submitted for genotyping as described above but conducted at the Institute de Biologie Intégrative et des Systèmes (IBIS) at Université Laval. At both facilities, target variants were scored using the Torrent Suite software VariantCaller plugin (ThermoFisher Scientific) using the original designed target SNP file (i.e., Cgig v.1.0 hotspots; see *Data Availability*). The custom pipeline *amplitools* (see *Data Availability*) was used to filter multi-locus genotypes files for target variants and convert to genepop format. Below, the population genetic samples and the VIU samples are referred to as the ‘pilot dataset’, and the MBP CHR8 samples are referred to as the ‘MBP CHR8 dataset’.

### Repeatability, filtering, and population genetic analyses

The pilot dataset genepop file was imported into R using adegenet and analyzed using the custom pipeline *simple_pop_stats* using instructions provided in *ms_oyster_panel* (see *Data Availability*). Technical replicate pairs with at least a 50% genotyping rate per individual were used to determine the proportion of concordant genotypes. Downstream analyses used the replicate individual with the highest genotyping rate. Samples, then loci, were filtered to keep those with less than 30% missing genotypes. Monomorphic loci, and loci that were previously determined to be align to more than one position in the genome (*see above*) were also removed. Although markers were characterized for deviation from Hardy-Weinberg proportions in each population using the *hw.test* of pegas, deviating markers were not removed, only flagged.

The dataset was converted to a genlight object using dartR (v.2.0.4; Gruber et al., 2018). Inter-individual relatedness between individuals was estimated using the Ritland statistic (Ritland, 1996) as implemented in related (v.1.0; Pew et al., 2015). First-degree relatedness values were estimated using known full-sib pairs in the VIU dataset (*see below*). Pairs of individuals within a population that had relatedness values greater than the lowest value for known first-degree relatives were subjected to a process of elimination where one individual per pair was removed until no pairs remained above the threshold. The purged first-degree relative dataset was then analyzed by principal components analysis (PCA) using the function *glPca* in adegenet, retaining PC1-4, and population averaged *F*_ST_ (Weir & Cockerham, 1984) was calculated for any population with more than 15 retained individuals using hierfstat (v.0.5-11; Goudet & Jombart, 2022). The full dataset with putative first-degree relatives included was used to identify private alleles in each genetic grouping using poppr (v.2.9.3; Kamvar et al., 2014), where Japan, France, and Canada (Pendrell) were pooled together, and the three VIU generations were all pooled together. The hatchery or farm collections were kept as separate groupings in this analysis. The full dataset was used to calculate per locus mean *F*_ST_ using the *Fst* function of pegas, and per locus heterozygosity using the *summary* function of adegenet. The data for the MBP CHR8 families was used as a test dataset and filtered as above, removing individuals then loci with more than 30% missing genotypes, and removing monomorphic loci. This dataset was used for parentage analysis to compare with the VIU parentage analysis in order to assess the utility of the panel in other programs (*see below*).

### Parentage assignment

The pilot dataset was reduced to only the VIU breeding program samples (now referred to as the VIU dataset), filtered again to keep loci with MAF > 0.01 and to remove monomorphic loci, then converted from genind format to rubias format (Moran & Anderson, 2019) using the function *genepop_to_rubias* of *simple_pop_stats* (see *Data Availability*). Data preparation and analysis for parentage closely followed vignettes of *close-kin mark recapture-sim* (CKMR-sim; Anderson, 2024). Using allele frequencies estimated from the VIU dataset, a simulated set of parent-offspring, full-sib, half-sib, and unrelated relationships were created to determine the power for the amplicon panel genotypes to resolve different relationship types, and to estimate expected log likelihood values for the different relationship categories. The dataset was checked for closely matching potential duplicate samples. Parent-offspring relationships and full-sibling relationships in offspring and parents were evaluated using VIU samples (i.e., VIU_F2 vs. VIU_F1). Siblings of true parents were included in the analysis to test the power of the amplicon panel to resolve relationships in the present of closely related possible parents (e.g., ‘aunts’ or ‘uncles’).

Following analysis and grouping of individuals into putative families, pedigree information was provided and compared against the genetic parentage analysis relationships. Offspring that failed to assign to genotyped parents were tallied to assess false negatives, and incorrect assignments were tallied as false positives. The number of putative parents was not limited to two, but rather all assignments over the log likelihood cutoff (*logl* > 5) were retained in the results. This cutoff was based on the total number of comparisons and expected false positive rates as per the CKMR-sim tutorials. The percentage of the total possible assignments that could have been made was calculated for each family in both parentage analyses, and the percentage of the total assignments made that were false positives was also calculated. Suspected pedigree or handling errors were inspected alongside genetic sibship analyses and breeding records to reconcile conflicting relationship information. These errors, when resolvable, were corrected.

Following the initial analysis described above, empirically identified trios (i.e., offspring and two parents) with strong support were identified (i.e., both parents assigned with log likelihood values ≥ 10, only two parental assignments with log likelihood scores above the set cutoff). When a set of parents were supported by more than one trio, the trios were kept as the empirical trio results to be used to identify unexpected genotypes potentially indicating null alleles. The empirical trios were used to infer expected genotypes of offspring per locus and family, and the observed offspring genotypes were compared to expected offspring genotypes to tally the number of unexpected genotypes per locus. Finally, the parentage analysis with all samples was re-done as described above but with the following filtered datasets: (1) all loci passing filters; (2) loci with fewer than four offspring with incompatible genotypes. Percent completeness of assignments, and percent of assignments that were false positives were calculated for each of the above three parentage approaches. This same approach was also carried out independently with the MBP CHR8 dataset, and the problematic loci identified with each dataset were compared against each other to determine whether the VIU and the MBP CHR8 dataset had the same loci that were exhibiting potential null alleles or otherwise unexpected offspring genotypes.

## Results

### SNP selection for marker design and chromosomal positions

The input ddRAD-seq dataset (Sutherland et al., 2020) contained 366 Pacific oyster samples genotyped at 16,492 SNPs (single SNP retained per RAD-tag). It had been filtered to retain SNPs that were present in at least 70% of individuals in all 15 populations, with global minor allele frequency (MAF) ≥ 0.01, and not significantly out of Hardy-Weinberg equilibrium. The dataset was then limited to naturalized British Columbia (BC) samples, retaining 122 individuals from four populations. The BC-only dataset was filtered to keep SNPs with MAF ≥ 0.01, retaining 13,438 SNPs. Marker design candidates were selected for elevated heterozygosity (n = 300; *H*_OBS_ = 0.41-0.5) or elevated *F*_ST_ between naturalized populations within BC (n = 300; *F*_ST_ = 0.046-0.178), where nine markers were common to both lists. An additional 20 markers were selected due to being previously identified as private alleles (see *Methods*; Figure S1; Additional File S2). Of the 611 submitted SNPs for design, 19 were excluded, and 592 were designed into the panel (i.e., *Cgig_v.1.0*; see Additional File S3 for marker names, contig identifiers, positions and alleles).

Of the 592 design sequences, 499 (84.2%) aligned using bowtie2 to a single location in the chromosome assembly, 55 aligned to multiple locations, 12 aligned singly to non-chromosome scaffolds, and 45 did not map. However, when aligned with bwa mem, 25 of the design sequences aligned to multiple locations, and these were different than the bowtie2 putative multi-mappers (see locus identifiers in Additional File S4). Due to this uncertainty, the bowtie2 putative multi-mappers were removed from the present analysis, but remain in the targets file. Each chromosome had an average (± s.d.) of 49.9 ± 26.3 single-mapping amplicons per chromosome (range: 15-83; Figure 1). Several significant gaps in chromosome coverage with single mapping markers were observed (Figure 1).

**Figure 1.**
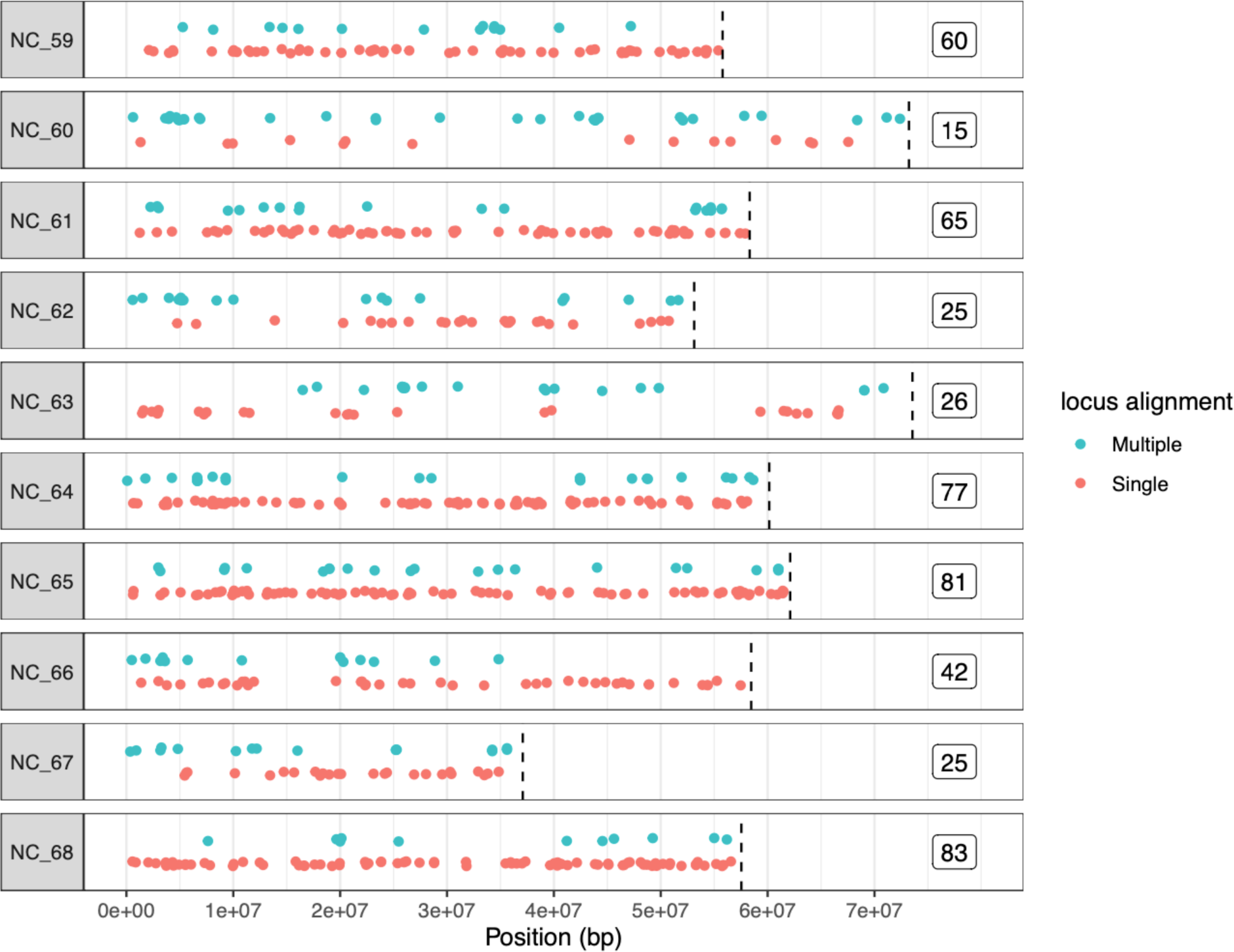
Positions of design sequences for the amplicon panel in the chromosome-level reference genome (GCF_902806645.1; Peñaloza et al., 2021). Most design sequences mapped to a single location (n = 499; red/lower), but some mapped to more than one position (teal green/upper, showing multiple locations for single amplicons), and were subsequently removed from the analysis. Chromosome ends are shown by hatched vertical lines, and the number of singly mapping design sequences per chromosome are shown to the right of the plot. Abbreviated NCBI chromosome names are shown to the left of the plot (e.g., NC_047559.1 to NC_59).

### Panel genotyping and repeatability

The pilot dataset comprised 543 samples including technical replicates and controls, or 370 unique samples and 32 negative controls. Samples were genotyped at 592 amplicons across two sequencing runs (i.e., two sequencing chips). Seven amplicons were uncalled in both chips, leaving 585 genotyped amplicons remaining. The median of the average amplicon depth per individual was 172.5x and 180.1x for the two chips, whereas negative controls were 0.90x and 5.3x, respectively (Figure S2). On the second chip, three of the 19 negative control wells had mean marker coverage greater than 40x, suggesting some contamination in these wells.

Technical replicate sample pairs (n = 68 pairs) showed average (± s.d.) genotype concordance of 97.5 ± 4.2% for SNPs genotyped in both samples (mean = 460 SNPs genotyped in both samples of the pair, with a total of 31,303 genotypes). The most frequent discordant genotypes between technical replicates were homozygous alternate and heterozygous genotypes (n = 552; 1.76%), and less frequent were homozygous reference and heterozygous genotypes (n = 153; 0.49%) or least frequent as homozygous reference and homozygous alternate genotypes (n = 4; 0.01%). Ten replicate sample pairs had more than 5% of the markers discordantly called; on average, these pairs had 359.5 loci typed in both samples, whereas the remainder pairs on average had 477.7 loci. One replicate pair had genotype discordance above 10%, and this pair was only genotyped at 342 loci. Therefore, samples with lower genotyping rates also had lower concordance.

### Filtering and population genetics

The 370 oysters in the pilot dataset were comprised of 181 oysters from the VIU breeding program, 71 from aquaculture programs (i.e., hatcheries or farms), and 63 and 55 from naturalized and wild sources, respectively (Table 1). After filtering samples for genotyping rate (GR > 70%), 312 samples were retained (Table 1; Figure S3). Filtering loci for genotyping rate (GR > 70%) removed 111 loci (Figure S4) leaving 474 loci remaining. Removing monomorphic loci removed an additional 43 loci, resulting in the retention of 431 loci. Any remaining of the 55 multi-mapping loci as determined by the bowtie2 alignment were also removed at this stage, dropping 22 additional loci and retaining 409 loci. Retained markers showed a bimodal *H*_OBS_ distribution (Figure 2A). The upper *H*_OBS_ grouping (*H*_OBS_ > 0.25; n = 246) has 221 loci that were included in the panel due to high *H*_OBS_. The lower *H*_OBS_ grouping (*H*_OBS_ < 0.25; n = 163) is mainly comprised of loci included due to high *F*_ST_ (i.e., n = 132), as well as those included based on presence as private alleles (n = 11). Per locus *H*_OBS_ and *F*_ST_ values for the 409 retained loci are shown in Additional File S5. Loci Hardy-Weinberg proportions (HWP) were inspected for wild (JPN, CHN) and naturalized populations (PEN, FRA) finding 15, 24, 26, and 26 variants out of HWP, respectively. This includes 11 loci that were out of HWP in two populations, and one marker in three populations. These markers were not removed from the analysis but can be referenced for future consideration in Additional File S6.

**Figure 2.**
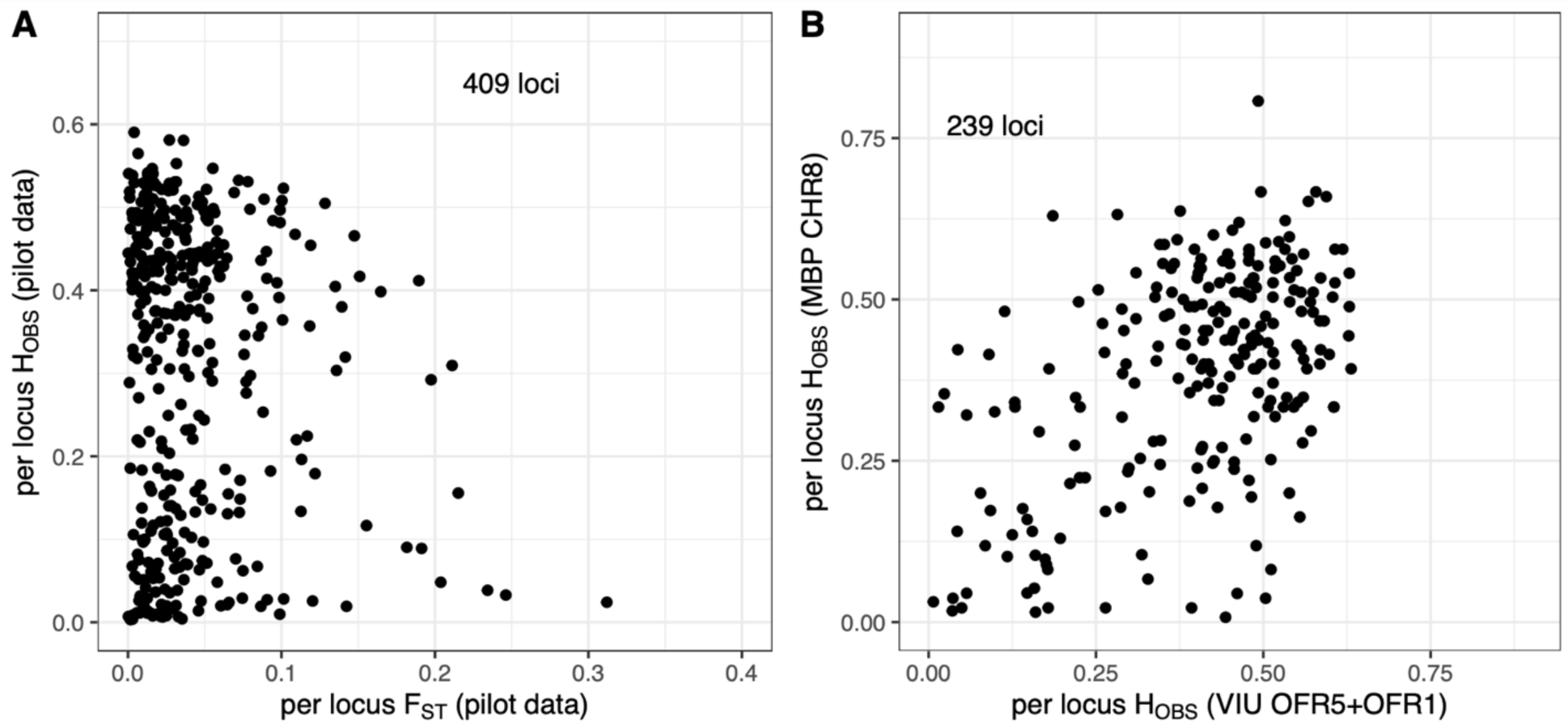
(A) Per locus observed heterozygosity (*H*_OBS_) vs. genetic differentiation (*F*_ST_) in the pilot study dataset for all filtered loci. (B) Per locus *H*_OBS_ for the MBP CHR8 parentage dataset vs. per locus *H*_OBS_ for the VIU OFR5 and OFR1 parentage dataset for all retained loci in both datasets.

The filtered dataset (n = 312 individuals, 409 loci) was then inspected for highly related pairs of individuals within each population (Figure S5). Outlier pairs with elevated relatedness were observed in most populations. The known full-sibs in the largest cross event in the VIU population used (i.e., oyster family run 5; OFR5) were used to calculate average relatedness values within each family, and the 14 families had an average pairwise relatedness value with the Ritland statistic of 0.43, and the average minimum relatedness value per family was 0.26, which was used as a cutoff to remove highly related individuals specifically to reduce family effects on the PCA and between population *F*_ST_. This resulted in the removal of most of samples from the hatchery or farm populations (i.e., VIU, DPB, and GUR) for PCA and *F*_ST_ analyses. Populations with fewer than 15 individuals were not used for *F*_ST_ calculation.

A PCA with putative first-degree relatives removed found general overlap of all populations except CHN, which were separated from the rest of the samples across PC1 (3.2% variance explained) and PC2 (2.2% variance explained), respectively (Figure S6). Similarly, the Japan translocation lineage (i.e., JPN, PEN, FRA) had low *F*_ST_, with the lower range of the 95% confidence intervals overlapping zero (i.e., no differentiation) and the upper range reaching *F*_ST_ = 0.012 (Table S2). By contrast, CHN had higher differentiation, for example JPN-CHN 95% C.I. *F*_ST_ = 0.023-0.048).

Private alleles were empirically identified in the genotyped dataset (with putative first-degree relatives included) at the regional level (i.e., JPN lineage, VIU, DPB, GUR, CHN). In total, 20 private alleles were observed, including some with high frequency. Three markers were included in the panel because they contained private alleles in DPB (Sutherland et al., 2020); although different samples were used here, two of these loci remained after filters and also were observed to be private alleles in this dataset, including locus 39139 (eight occurrences), and locus 538043 (six occurrences). Similarly, locus 336452, included in the panel as a private allele within GUR in Sutherland et al. (2020) was also private in this dataset (nine occurrences). One high frequency private allele was observed in the VIU samples with eight observations (locus 590965). However, this marker would have been designed based on presence in other BC samples, and so is likely just a rare allele that has been enriched in the VIU broodstock program. A private allele was also observed for each of CHN and JPN with five observations (see full list in Table S3).

### Parentage assignment in the VIU dataset

The panel was tested on families from the oyster family run 5 (OFR5) spawning event of the Vancouver Island University (VIU) Pacific oyster breeding program. OFR5 involved 15 crosses using 18 unique parents that originated from cross events OFR1 and OFR2, including nine dams and nine sires, all of which were genotyped. One OFR5 broodstock parent, BR22, was missing. In total, 91 OFR5 offspring (81.3%) were retained after filters (Table 1, Table 2).

**Table 2.** Parentage analysis of VIU OFR5 using filtered data (including incompatible genotype removal, see Methods; n = 328 loci included). In total, 160 of 172 potential assignments were made (93.0% complete), and eight false positives occurred (4.8% of calls). Per family, the individual identifier (with ‘BR’ prefix) for dam and sire are shown alongside their originating family identifier (brackets) and their number of retained genotyped markers (parentheses). Per family, the number of genotyped offspring is shown with the number of correct two- or one-parent assignments, the percent completeness of the assignments (% ID), and any incorrect assignments. All assignments above the applied cutoff were retained (see Methods). Acronyms: pat. = paternal; mat. = maternal

| OFR5 family | Dam [fam.] (loci) | Sire [fam.] (loci) | Off-spring | Two-parent assign. | One-parent assign. | % ID | Incorr. assign. |
| --- | --- | --- | --- | --- | --- | --- | --- |
| OFR5_2 | BR19 [2-6] (323) | BR11 [1-14] (326) | 6 | 6 | - | 100.0 |  |
| OFR5_3 | BR14 [2-10] (306) | BR2 [1-12] (322) | 9 | 8 | 1 | 94.4 |  |
| OFR5_4 | BR14 [2-10] (306) | BR11 [1-14] (326) | 6 | 6 | - | 100.0 |  |
| OFR5_5 | BR3 [1-4] (317) | BR10 [1-18] (325) | 6 | 5 | 1 | 91.7 | 1 x pat. sib |
| OFR5_7 | BR17 [2-11] (323) | BR4 [1-10] (317) | 7 | 7 | - | 100.0 |  |
| OFR5_8 | BR3 [1-4] (317) | BR7 [2-11] (316) | 8 | 7 | 1 | 93.8 |  |
| OFR5_9 | BR9 [1-16] (318) | BR23 [2-6] (324) | 8 | 8 | - | 100.0 |  |
| OFR5_10 | BR25 [1-19] (322) | BR23 [2-6] (324) | 7 | 5 | 2 | 85.7 | 3 x mat. sib |
| OFR5_12 | BR14 [2-10] (306) | BR20 [1-19] (326) | 7 | 7 | - | 100.0 | 4 x pat. sib |
| OFR5_13 | BR22 [2-14] (0) | BR26 [1-9] (325) | 3 | na | 3 | 100.0 |  |
| OFR5_14 | BR24 [1-13] (318) | BR2 [1-12] (322) | 8 | 8 | - | 100.0 |  |
| OFR5_15 | BR18 [2-13] (320) | BR10 [1-18] (325) | 2 | 2 | - | 100.0 | 1 x pat. sib |
| OFR5_16 | BR22 [2-14] (0) | BR2 [1-12] (322) | 7 | na | 4 | 57.1 |  |
| OFR5_17 | BR21 [1-1] (322) | BR5 [2-21] (319) | 8 | 5 | 2 | 75.0 |  |

An initial analysis of the data found likely pedigree or handling errors. Four samples were identified that were suspected handling errors in which the offspring had the incorrect family label. These were evident using parent-offspring, or sibship analyses. Second, family OFR5-15 was found to be a mixed family, with two separate sets of parents (i.e., the expected parents BR18 and BR10, or the parents of OFR5-14, BR24 and BR2; Table 2). Third, a pedigree error was identified where the sire of family OFR5-2 was BR11 and not BR2 (family 1-12). These were all corrected before subsequent analyses.

To identify potential null alleles in the amplicon loci, reliable two-parent assignments were identified and used to identify incompatible offspring genotypes based on the observed parental genotypes (see Methods). Reliable assignments were defined as those trios with exactly two parents assigned above an elevated cutoff. Putative families were estimated from the reliable assignments, keeping parental pairs as a putative family when more than one observation of a parental pair occurred. These identified trios included 37 offspring from 11 families. Within these trios, 36 loci were observed with at least four offspring with incompatible genotypes, and 95 loci with at least two offspring with incompatible genotypes (Figure S7). Parentage datasets were constructed to evaluate the impact of these loci exhibiting potential null alleles: (i) keeping all filtered loci (n = 364 loci retained); or (ii) removing loci with four or more incompatible genotypes (n = 328 loci retained). An improvement in assignments occurred and a reduction of false positives by removing loci with at least four incompatible offspring genotypes, and so this approach was taken (Table 3). In this dataset, the OFR5 broodstock was genotyped with an average of 313.2 loci per individual (Table 2), and the offspring at an average of 307.0 loci per individual (Additional File S7).

**Table 3.** Comparative parentage assignment for OFR5 and MBP CHR8 families with all filtered loci or filtered loci with those with at least four incompatibilities observed removed. The number of loci present in the dataset, the number of assignments and percentage of possible assignments, as well as the number of false positives and percentage of assignments made as false positives are shown. The datasets with loci having more than four incompatibilities removed were used for downstream analyses.

| Cross event/<br>families | Locus filter | Num. loci | Num. assign. | Num. possible assign. | Percent assigned | Num. false positive (FP) | Percent assigns as FP |
| --- | --- | --- | --- | --- | --- | --- | --- |
| OFR5 | all filtered loci | 364 | 134 | 172 | 77.9% | 11 | 7.6% |
| OFR5 | Remove loci with ≥ 4 incompatibilities | 328 | 160 | 172 | 93.0% | 8 | 4.8% |
| MBP CHR8 | all filtered loci | 324 | 163 | 215 | 75.8% | 7 | 4.1% |
| MBP CHR8 | Remove loci with ≥ 4 incompatibilities | 289 | 185 | 215 | 86.0% | 17 | 8.4% |

Within the 328-loci dataset, 160 of 172 potential assignments were made (93.0% complete), and eight false positives occurred (4.8% of assignments made; Table 3; Table 2). The greatest number of false positives occurred in families F10 and F12, where maternal or paternal siblings were assigned in addition to the correct assignment. All parent-offspring relationships are shown in Figure 3, and assignments with accompanying log likelihood ratios are shown in Additional File S8.

**Figure 3.**
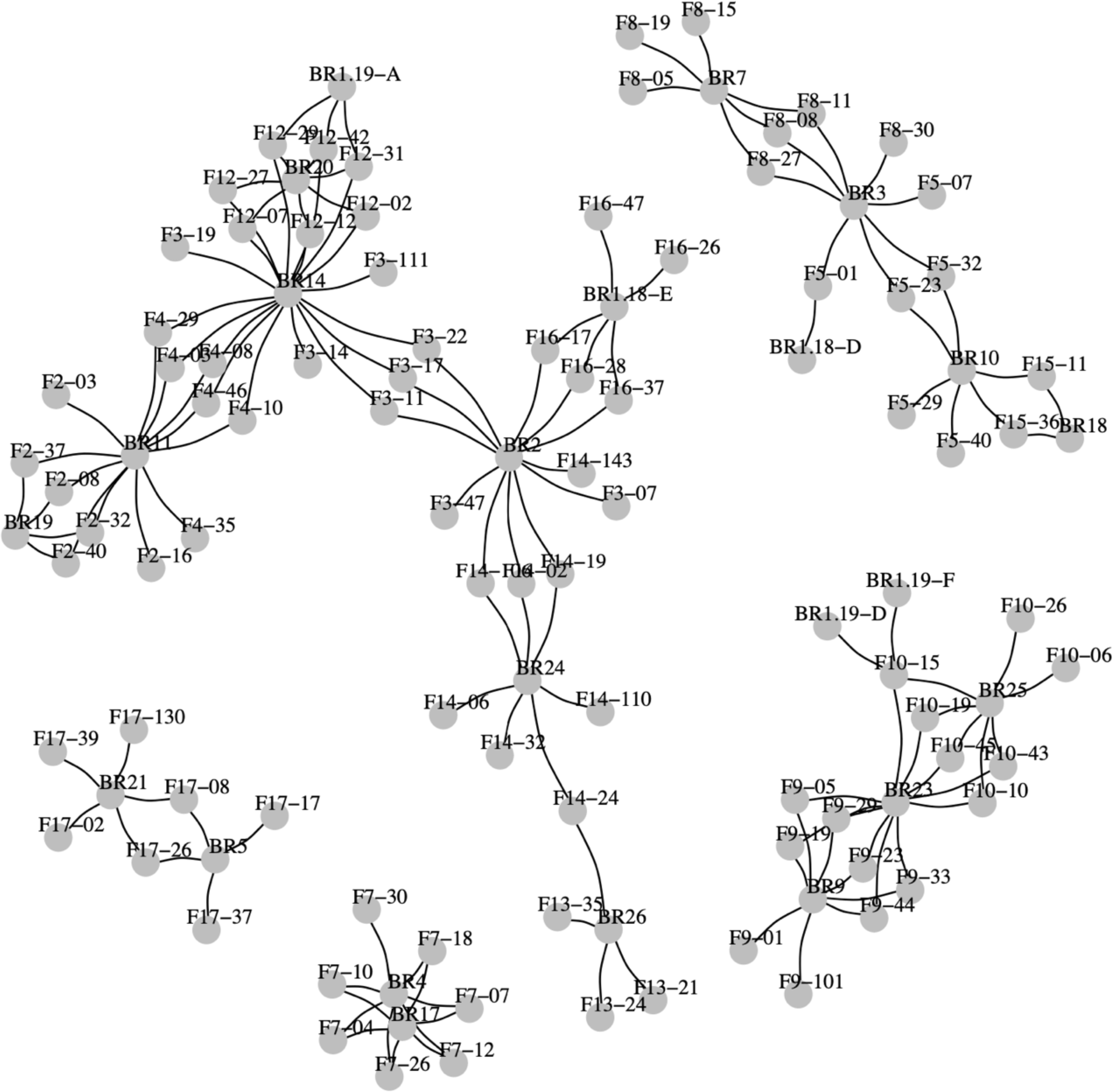
Parent/offspring (PO) relationship network for the VIU OFR5 cross showing all putative PO relationships with log likelihood > 5 using 328 SNP loci. The offspring are denoted with “F” (e.g., F17-37) and the potential parents with “BR” (e.g., BR9). For completeness, all putative linkages are shown, including more than two per offspring individual.

### Parentage assignment in the MBP CHR8 dataset

An independent dataset comprised of families from the Molluscan Broodstock Program (MBP CHR8) was genotyped to further evaluate the panel performance for parentage assignment. This dataset was expected to be more challenging for the panel given the longer captive breeding history and higher relatedness among broodstock. The MBP spawn was comprised of 16 families from MBP cohort 27 (seventh generation of selective breeding) including 25 unique broodstock (nine dams and 16 sires). All parents were genotyped, but one parent (65_8F) was removed due to low genotyping rate. In total, 111 offspring (100.0%) were retained after filtering (Table 1, Table 4). The dataset was comprised of 576 scored loci, but after filtering the excess missing data (missing ≥ 30%), 519 loci were retained. An additional 174 monomorphic loci were removed, leaving 345 loci. Removing the multi-mapping loci from the bowtie2 analysis resulted in an additional 21 loci being removed, leaving 324 loci remaining. The MBP CHR8 family distributions of the likelihoods of simulated relationships were similar to those of the VIU OFR5 families (Figure S8). Further, the observed heterozygosity per locus of the VIU OFR5 family and MBP CHR8 shows many loci with similarly elevated H_OBS_ in both datasets (Figure 2B).

**Table 4.** Parentage analysis of MBP CHR8 families using filtered data (including incompatible genotype removal, see Methods; n = 289 loci included). In total, 185 of 215 potential assignments were made (86.0% complete), and 17 false positives occurred (8.4% of calls). See VIU OFR5 table for full table explanations.

| CHR8 family | Dam [fam.] (loci) | Sire [fam.] (loci) | Off-spring | Two-parent assign. | One-parent assign. | % ID | Incorr. assign. |
| --- | --- | --- | --- | --- | --- | --- | --- |
| 101 | 55_3F [55] (281) | 55_16M [55] (284) | 7 | 7 | - | 100.0 | 2 x [55] aunts |
| 102 | 55_3F [55] (281) | 55_14M [55] (284) | 7 | 7 | - | 100.0 |  |
| 103 | 55_45F [55] (283) | 55_17M [55] (285) | 7 | 7 | - | 100.0 | 6 x [55] uncles |
| 105 | 55_27F [55] (286) | 58_29M [58] (287) | 6 | 6 | - | 100.0 | 1 x [58] uncle,<br>1 x [55] aunt |
| 106 | 55_27F [55] (286) | 58_20M [58] (284) | 7 | 7 | - | 100.0 |  |
| 107 | 55_45F [55] (283) | 58_6M [58] (287) | 7 | 1 | 6 | 57.1 | 3 x [58] uncle,<br>1 x [58] aunt |
| 108 | 55_5F [55] (286) | 65_5M [65] (281) | 7 | 2 | 5 | 64.3 |  |
| 109 | 55_5F [55] (286) | 65_3M [65] (286) | 7 | 4 | 3 | 78.6 |  |
| 110 | 55_45F [55] (283) | 65_11M [65] (286) | 7 | 4 | 3 | 78.6 |  |
| 111 | 55_2F [55] (255) | 79_6M [79] (286) | 6 | 4 | 2 | 83.3 | 2 x [58] unkn. |
| 112 | 55_2F [55] (255) | 79_4M [79] (281) | 7 | 2 | 5 | 64.3 |  |
| 113 | 55_45F [55] (283) | 79_18M [79] (284) | 7 | 6 | 1 | 92.9 |  |
| 114 | 65_4F [65] (286) | 58_33M [58] (287) | 7 | 7 | - | 100.0 |  |
| 115 | 65_8F [65] (0) | 79_13M [79] (282) | 7 | na | 7 | 100.0 | 1 x [65] uncle |
| 116 | 55_41F [55] (287) | 65_19M [65] (282) | 7 | 4 | 3 | 78.6 |  |
| 117 | 58_9F [58] (287) | 79_1M [79] (284) | 7 | 7 | - | 100.0 | 1 x [58] uncle |

An initial analysis found suspected handling errors, including three offspring with incorrect family identifiers. Offspring were labeled as family/offspring number. Incorrect offspring identified were as follows: 102/8 was from family 106 and not 102; 106/1 was from family 113 and not 106; and 113/8 was from family 102 and not 113. These were sequential in DNA extraction and library preparation, and so the series of unexpected assignments likely reflects a handling error in DNA extraction or library preparation and not in sampling. These errors were corrected by providing new identifiers to the unexpected offspring to match their correct families, and then the same approach to identify putative null alleles was taken as described above for OFR5 (see above and Methods). Empirically determined trios with strong evidence were identified in the MBP CHR8 families, including 38 offspring from nine different families. Using these trios, 73 loci were identified that had at least two offspring with incompatible genotypes based on expected genotypes inferred from parents, and 35 loci with at least four offspring with incompatible genotypes. The two datasets (i.e., all filtered loci: 324 loci; remove when ≥ 4 incompatibilities: 289 loci), were used for parentage analysis. The dataset with 289 loci resulted in the most assignments, but also the most false positive assignments (Table 3, Table 4). In this dataset, the parent generation (potential broodstock) were genotyped on average at 283.4 loci per individual (Table 4), and the offspring on average at 284.5 loci per individual (Additional File S7). This dataset had 185 of 215 possible assignments (86.0% complete), and 17 false positives occurred (8.4% of assignments). All identified parent-offspring relationships are shown in Figure 4, and assignment details in Additional File S8.

**Figure 4.**
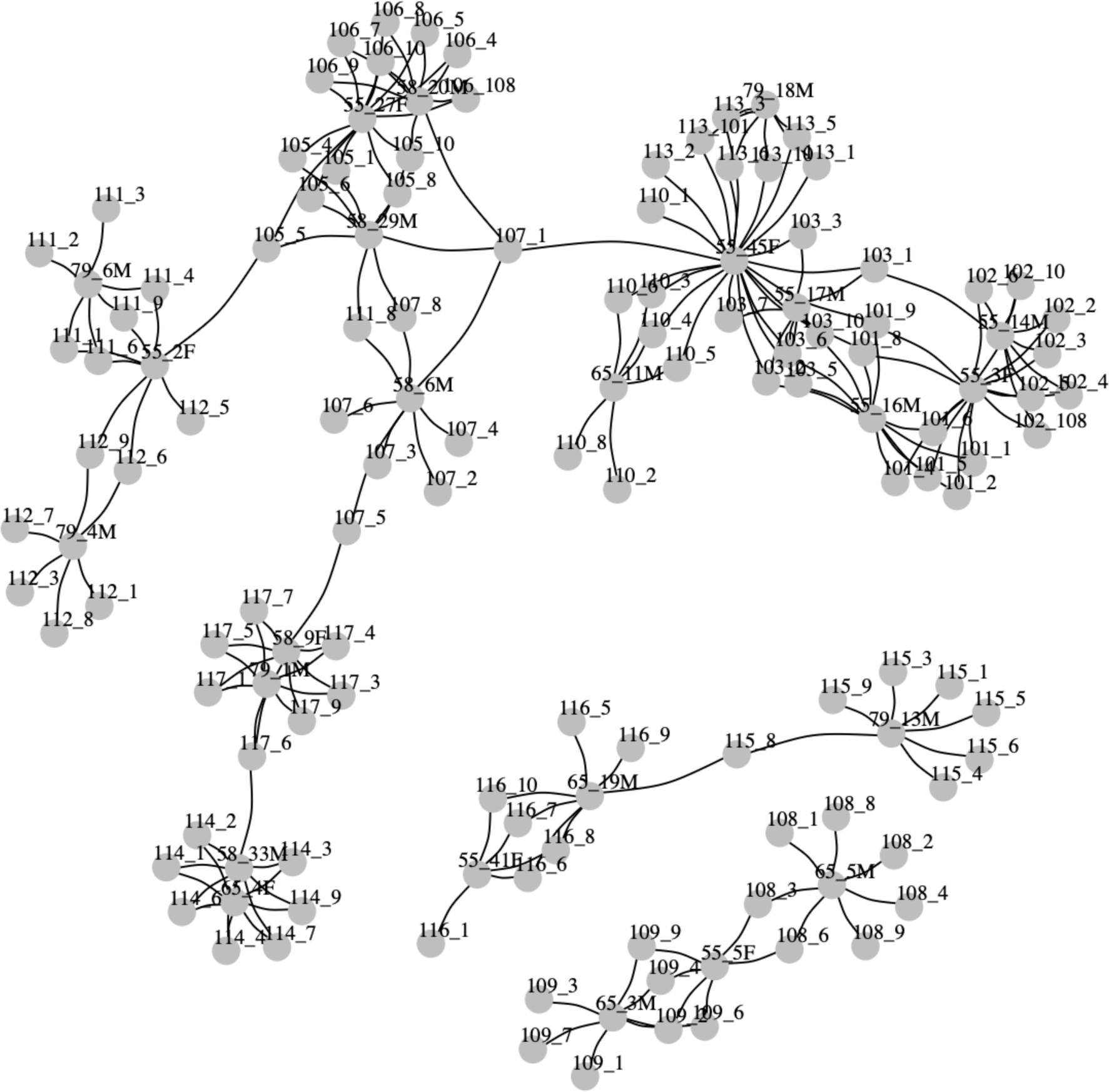
Parent/offspring (PO) relationship network for the MBP CHR8 families showing all putative PO relationships with log likelihood > 5 using 289 SNP loci. The offspring are shown with numeric family identifiers followed by an individual identifier (e.g., 108_4). The putative parents are also shown this way but also including a sex identifier (e.g., 65_5M). For completeness, all putative linkages are shown, including more than two per offspring individual.

To determine whether a common set of loci were exhibiting genetic incompatibilities (i.e., putative null alleles) in the VIU OFR5 and the MBP CHR8 families, overlap was inspected between the incompatible loci. Of the 36 loci that were incompatible in VIU (≥4 incompatibilities) and the 35 incompatibility loci in MBP, only seven were found common in both datasets (i.e., 19% or 20% of the total VIU or MBP loci, respectively). This low overlap occurred even though 35 of 36 of the incompatible VIU loci were present in the MBP CHR8 dataset (before incompatibles were removed), and 28 of 35 of the incompatible MBP CHR8 loci were present in the VIU dataset (before incompatibles were removed; Additional File S9). This suggests that putative null alleles identified are largely population specific.

## Discussion

This work demonstrates the effectiveness of the Pacific oyster Cgig_v.1.0 amplicon panel for characterizing genetic variation and conducting parentage assignment, as well as provides the analysis framework to support additional use of the panel, as well as future development of the panel in subsequent design iterations. The unique characteristics of the Pacific oyster, including high levels of genetic polymorphism, likely led to challenges faced in the analysis, such as the presence of numerous, population-specific null alleles. Initial results were sufficient to detect suspected handling or pedigree errors, including in breeding program samples from populations not part of the panel design, and with careful filtering steps, parentage assignment success was able to be improved. We expect that the tools provided here will bring advances to Pacific oyster breeding programs for both small and large operations.

### Maintaining genetic diversity and supporting breeding program development

Several genetic approaches may be used to increase the resiliency of cultured Pacific oysters. First, maintaining overall genetic diversity retains variation required for rapid adaptation (Barrett & Schluter, 2008) and avoids significant inbreeding depression that can impact growth and survival (Evans et al., 2004; Wada & Komaru, 1994). Second, artificial selection methods including family-based selection (i.e., breeding families that show most favourable phenotypes as selection targets), marker-assisted selection (i.e., selecting broodstock based on genotypes at loci associated with a phenotype), and genomic selection (e.g., breeding based on multilocus genotype relationships derived from a training set of phenotyped individuals), can be used to improve phenotypic targets of breeding populations (Boudry et al., 2021). Genomic or marker-assisted selection cannot be undertaken without genotyping. Similarly, maintaining diversity or separation of distinct breeding lines benefits from the ability to empirically measure diversity and confirm family relationships.

Maintaining genetic diversity in Pacific oyster breeding populations is important as rapid reductions in genetic diversity can occur (Hedgecock & Sly, 1990), as can small sustained reductions that accumulate over time (Carlsson et al., 2006; Varney & Wilbur, 2020; Xiao et al., 2011). Using genetic markers, the effective population size and other measures of diversity such as allelic richness can be directly monitored in the breeding population (Gurney-Smith et al., 2017; Hedgecock & Sly, 1990). Allelic richness may be a more appropriate measure to monitor diversity than observed heterozygosity or per-individual inbreeding coefficients (Sutherland et al., 2023; Varney & Wilbur, 2020; Yu & Guo, 2004), in part because these other metrics can be skewed by crossing divergent genetic lines for heterosis, an approach sometimes applied within the bivalve aquaculture industry. When reduced diversity is detected, solutions may be needed to replenish diversity (Li et al., 2007). Some countries, including Canada, have naturalized populations of conspecifics that can be incorporated within breeding programs (Sutherland et al., 2020) to increase diversity and avoid inbreeding. Having tools to empirically measure genetic diversity is important for breeding programs (Gurney-Smith et al., 2017) for a range of industry scales (Boudry et al., 2021).

Family-based selection or crossing of divergent lines both require maintenance of separate genetic groups, and benefit from confirming crosses and pedigree information to correct errors that may occur during animal husbandry or data recording. This has been a subject of concern for many decades (Hedgecock & Davis, 2007), and has led to the development of high-resolution melt curve genotyping panels (Sun et al., 2015; Zhong et al., 2013). Pedigree errors have been identified in the present work and elsewhere (Hedgecock & Pan, 2021) that would not have been identified without the use of a low-cost, effective genotyping panel, and if or when left unchecked, could slow progress in family-based breeding programs (Hedgecock & Pan, 2021). The amplicon panel presented here provides a rapid and lower-cost tool to evaluate the parentage of crosses and correct pedigree errors, thereby increasing the precision of family-based selection protocols.

There are many potential sources of error that can disrupt maintenance of separate genetic groups and directional family-based selection. They may occur during spawning, larval rearing, or growth phases, and can be caused by incorrect labeling, contaminating families (i.e., inadvertent mixing), as well as handling and record keeping errors. Many of these errors disrupt pedigree relationships, and if left uncorrected, can compound over time and impede directional selection progress (Hedgecock & Davis, 2007). As presented here, we identified several different sources of error: (1) mislabeled single offspring; (2) a full-sib family contaminated with a second full-sib family (i.e., the preceding family); (3) handling or recording errors for the sire of two families within one spawn event; (4) handling errors after sampling during laboratory preparation. Importantly, as discussed by Hedgecock and Pan (2021), these errors could impede selective breeding gains were they not detected, and may increase the mismatch between the genetic breeding values and the phenotypic outcomes of trials.

The amplicon panel may bring additional benefits to aquaculture programs. Currently, the application of genomic selection in shellfish aquaculture is limited by high genotyping costs per individual (Delomas et al., 2023). One framework, described and recently evaluated *in silico* for Pacific oyster by Delomas et al. (2023), is to use whole-genome resequencing of parents, lower-density amplicon panel genotyping of offspring, and imputation to enable more cost-effective application of modern genomic selection methods. Furthermore, field deployments for phenotyping trials can be conducted in common garden replicates, rather than using separate nets that confound spatial and family effects, by using genetic parentage analysis enabled by lower cost genetic marker panels (Allen et al., 2020). Finally, future versions of the panels can be designed to include putatively functional markers for marker-assisted selection, for example for shell colour or body size, or traits which can increase survival from thermal stress (Raymond et al., 2022), or for field survival likely linked to disease resistance (e.g., OsHV-1; Divilov et al., 2023).

### Future directions

The current panel performed well on the populations included in this analysis. Future work should continue to evaluate the panel in other populations to determine any ascertainment bias and to assess how broadly applicable the tool can be. As an example, in the present study, moving to different populations that were not involved in design, such as MBP’s CHR8 families, resulted in a specific loss of SNPs selected due to high *F*_ST_ in BC, whereas the SNPs selected for high heterozygosity were more likely to pass filters. The high *F*_ST_ markers are the most likely candidates to remove in future panel iterations. In this project, most genotyped individuals originated from populations that were included in marker discovery and comprised oysters from the Japan translocation lineage, China, and several commercial aquaculture hatcheries (Sutherland et al., 2020). In new populations, other variants within the panel’s amplified sequences (i.e., not the target designed SNP) may prove more powerful for parentage, and these could be characterized via adjustments to the data analysis post sequencing, and not requiring any physical changes to the current panel. This is beneficial because physical changes to the primer composition would be more laborious and costly. This is a particular advantage for amplicon-based sequencing that is not possible in array or high-resolution melt curve genotyping.

Using non-target variants from the existing amplicons may provide improvements to the panel for parentage purposes without changes to the primer pool. This is apparent given that some markers within Cgig_v.1.0 in the pilot study had low heterozygosity and minor allele frequency, including those selected due to high differentiation potential or presence as private alleles. For technical replicates, low genotyping rate pairs had the lowest percent concordance, suggesting that poor genotyping rate is not only an issue for missing data, but the data that is present has a higher chance of miscalls. Therefore, emphasis should be placed on retaining high quality throughout all phases of tissue collection, library preparation, and sequencing to minimize the loss of data or the generation of poor quality data. End-users of the panel should determine levels of missing data that are allowable for their uses. As missing data was more prevalent in specific collections, it is likely that this is driven by DNA quality or quantity; other sample collections with similar genetic backgrounds to the high failure collections had more consistent and complete genotyping, suggesting that the missing data is not largely due to primer failures due to variants at binding sites but rather due to DNA quality or quantity. Including higher value markers (i.e., higher MAF) may improve resolution of secondary assignments to full sibs of the true parent (i.e., false positives) without a significant increase in false negatives (i.e., missing assignments). Increased statistical power to assign genetic relationships may also be realized through a microhaplotype approach (Baetscher et al., 2018), which is an active area of research for the current panel.

Other advances to the panel are possible but would require modifying the primer suite by adding new or removing existing amplicons. The uniformity of coverage across chromosomes could be improved, filling in gaps observed in the current work, and thereby improving the potential for genomic selection (Delomas et al., 2023). As discussed above, when additional variants with putatively functional roles (or close linkage with causal variants) are identified, such as the recently identified SNP on chromosome eight associated with increased field survival associated with OsHV-1 outbreaks (e.g., Divilov et al., 2023), these will likely be valuable to add to the panel for breeding purposes. Although rigorous documentation and standardization of locus names is required, the malleability of the amplicon panel is a major strength, as loci can be added or dropped by changing the primer composition of the primer pool. Once it can be more conclusively determined which loci are true multi-mapping loci, for example through detecting more than two alleles in diploid individuals with microhaplotypes, these loci will be permanently removed from the panel. Furthermore, removing loci that exhibit null alleles would strengthen the use of the panel, however, the observation of the two different breeding programs largely showing different loci that had putative null alleles negates the ability to screen for null allele loci across a broad geographic scale without including this in an initial panel development protocol. In addition, given the high mutation rate of Pacific oysters, new mutations are likely to regularly emerge that could disrupt primer binding sites, producing new null alleles occurrences. Thus, users of the current panel are strongly encouraged to screen for null alleles within the target population of interest as removing these loci improves genetic parentage assignment success, as was demonstrated in the present work.

The amplicon panel provides a new tool for emerging Pacific oyster selection programs, but it is also important to also consider the ease of which the data can be analyzed. Although the tab-delimited genotype data produced by the genotyping platform may be easier to handle for those with less bioinformatics experience than high volume sequence data, moving from this output to the formats required for parentage or population genetic analysis does require moderate skills in data manipulation, quality control, and data management. With this in mind, we developed *amplitools*, a repository that is designed to make the use of this panel more accessible, including bash and R scripts for converting from raw variant calling output from standard sequencer software to a standard format used by numerous genetic analysis programs with a wide range of analysis types being available.

Some aspects of documentation within breeding programs will support the use of *amplitools*, such as not including third generations when running parent-offspring analyses, given the potential for grandparents to be miss-assigned as parents. Including parental sex information may also be beneficial for analysis, although this has not yet been implemented in *amplitools*. Additionally, we developed the associated repository *amplitargets* to keep and version control target files for genotyping, as well as any notes regarding changes between versions of SNP target files. Although the primer panel, *amplitools* and *amplitargets* are currently only implemented for the AgriSeq platform (ThermoFisher Scientific), the source marker sequences and target variants are all provided here and so they could be used for future panel development on other sequencing platforms, and the code could also be adapted and expanded to other data processing platforms.

## Conclusions

The Pacific oyster Cgig_v.1.0 amplicon panel is an effective tool for characterizing genetic variation, genetic similarity among populations, and relatedness between individuals, including for genetic parentage analysis applications. Combined with the *amplitools* workflow provided here, this amplicon SNP panel could benefit oyster breeding programs by facilitating monitoring of genetic variation and confirming pedigree relationships. The amplicon panel is a useful and lower-cost genotyping tool that has the ability to easily evolve and improve in future iterations by targeting novel variants and microhaplotypes *in silico*, as well as by more substantial revisions, such as including putatively adaptive loci, and achieving more uniform coverage across the genome.

## Supporting information

Supplemental Results

## Acknowledgements

Thanks to Chen Yin Walker (Vancouver Island University), Claire Rycroft (University of British Columbia), and the Molecular Genetics Lab of Fisheries and Oceans Canada for DNA extraction for the study. Thanks to those who collected or provided samples for this project, including Dr. Kristi Miller (DFO), Rob Saunders (RKS Labs), Prof. Nicolas Bierne and Dr. Ismaël Bernard (Eurêka Modélisation), Prof. Li Li and Dr. Sheng Liu, David Howell, J.P. Hastey (Nova Harvest Ltd.), Keith Reid (Odyssey Shellfish), Brian Yip (Fanny Bay Oysters), and Bruce Evans. Thanks to Geneviève Morin and Brian Boyle of IBIS (Université Laval) for support in implementing the amplicon panel for genotyping. This work was funded by the Gordon and Betty Moore Foundation (GBMF#5600) to CAS. This research used resources provided by the SCINet project of the USDA Agricultural Research Service, ARS project number 0500-00093-001-00-D. Funding for NFT was provided by USDA ARS CRIS project 2076-63000-005-000-D. Mention of trade names or commercial products in this publication is solely for the purpose of providing specific information and does not imply recommendation or endorsement by the U.S. Department of Agriculture.

## Data Availability

The amplicon marker panel referred to here as Cgig_v.1.0 is available commercially from ThermoFisher Scientific as SKU A58237 AGRISEQ PACIFIC OYSTER PANEL. The input data required for marker selection (single SNP per locus; plink and VCF formats), the VariantCaller multilocus genotypes for the pilot study, alignment results of submitted sequences against the reference genome, and sample population annotation is available on FigShare, DOI: 10.6084/m9.figshare.23646471

Three repositories support this project:

Manuscript code repository: https://github.com/bensutherland/ms_oyster_panel

*amplitools*: https://github.com/bensutherland/amplitools

*amplitargets*: https://github.com/bensutherland/amplitargets

