## Supplemental Results for "An amplicon panel for high-throughput and low-cost genotyping of Pacific oyster"

**Table of Contents:**


| | |
|---|---|
| **Figure S1. SNP selection for panel** | **Page 2** |
| **Figure S2. Marker sequencing depth** | **Page 3** |
| **Figure S3. Genotyping rate (samples)** | **Page 4** |
| **Figure S4. Genotyping rate (loci)** | **Page 4** |
| **Figure S5. Estimated relatedness** | **Page 5** |
| **Figure S6. Sample PCA** | **Page 6** |
| **Figure S7. Unexpected genotypes** | **Page 7** |
| **Figure S8. Simulated likelihoods** | **Page 8** |
| **Table S1. Selected loci in panel** | **Page 9** |
| **Table S2. Pilot study FST** | **Page 9** |
| **Table S3. Private alleles** | **Page 10** |

**A**

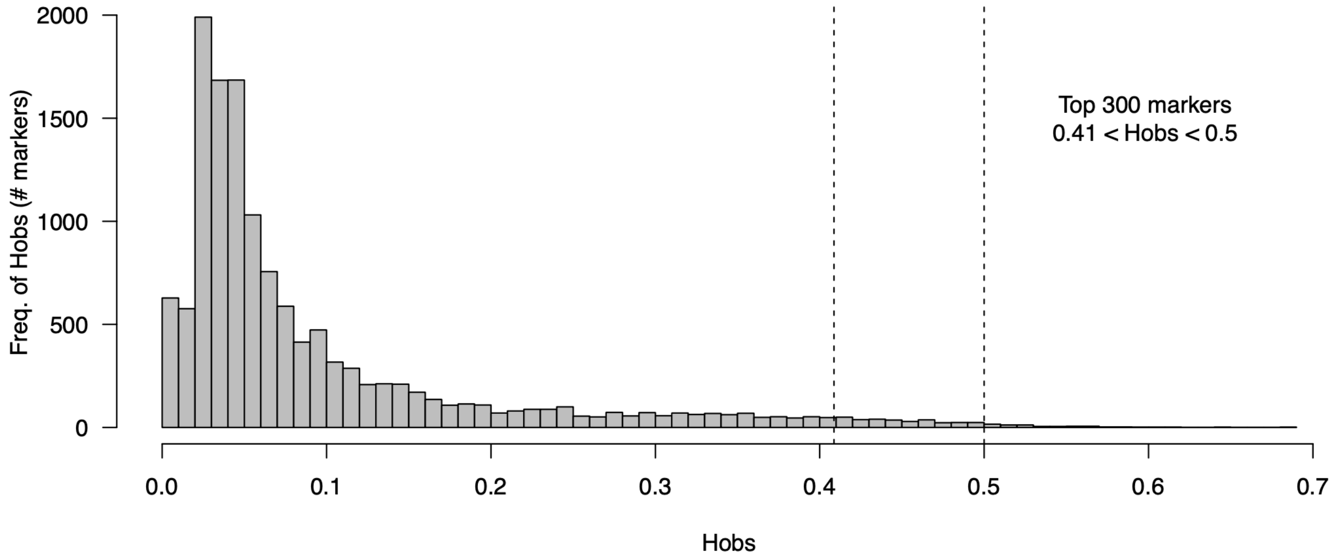


**B**

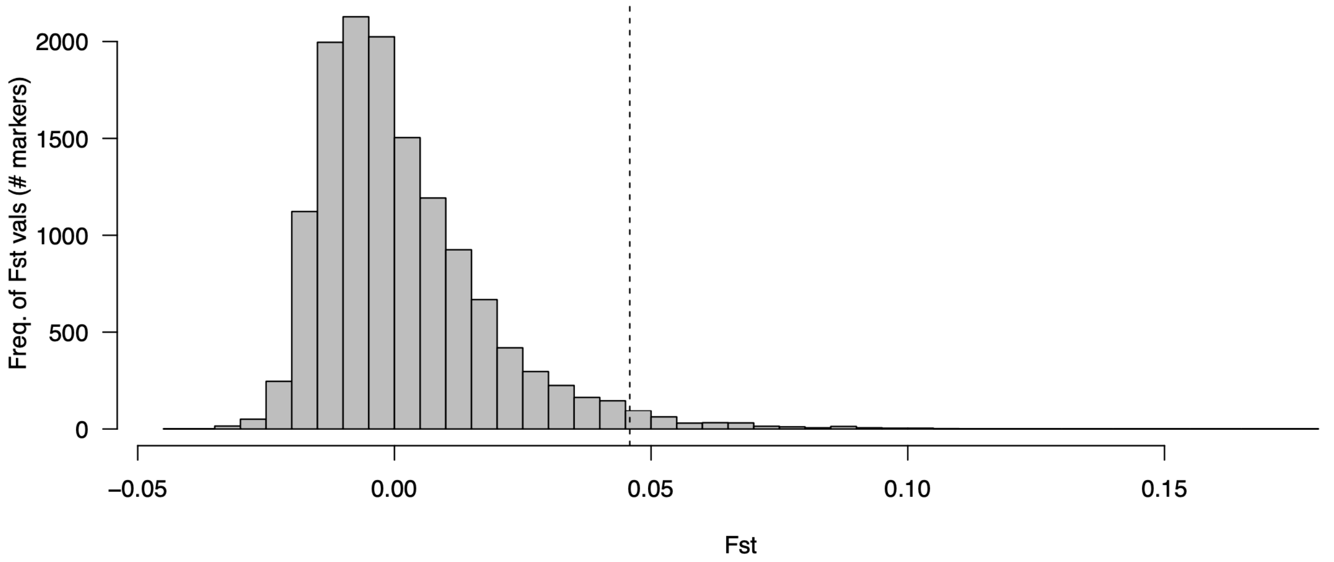


**Figure S1.** SNPs were selected to design into the marker panel using available ddRAD-seq data for naturalized oysters from British Columbia, Canada. SNP selection occurred by identifying the 300 markers with (A) highest observed heterozygosity below 0.5 (i.e., 0.41 < $H_{OBS}$ < 0.5), or with (B) highest genetic differentiation (i.e., $F_{ST} \geq 0.046$).

**(A)**

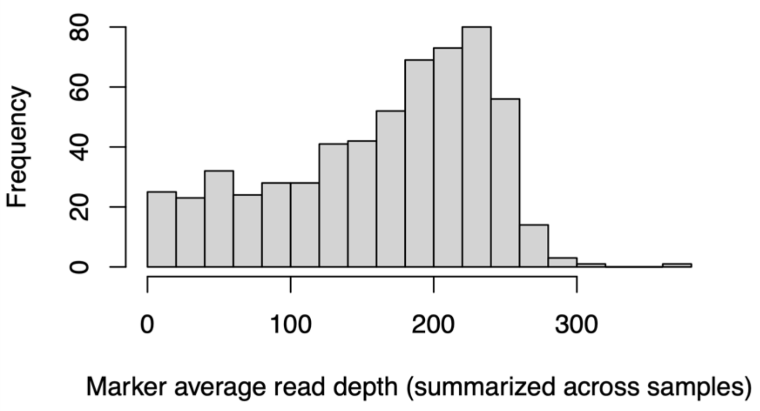


**(B)**

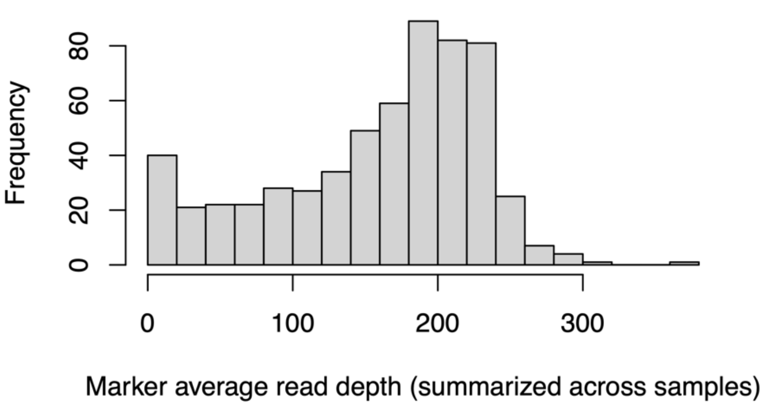


**(C)**

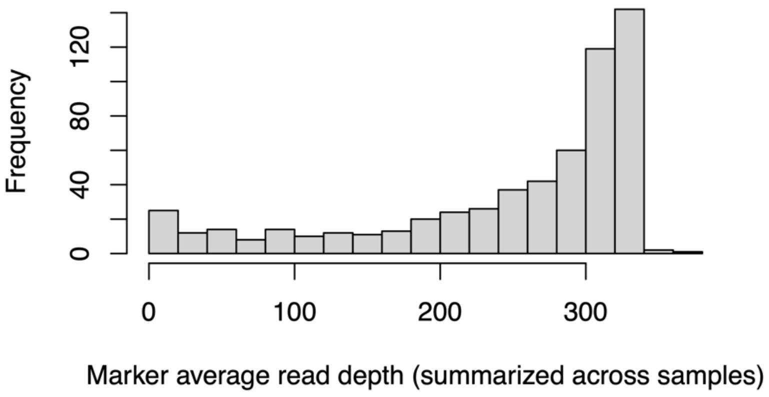


**Figure S2.** Frequency of per locus mean read depth for sequencing chips used in the pilot study (A-B; chips: R_2022_08_04_S5XL and R_2022_10_07_S5XL; or in the MBP CHR8 study (C); chip: R_2023_07_26_12_44_23_user_GSS5PR.

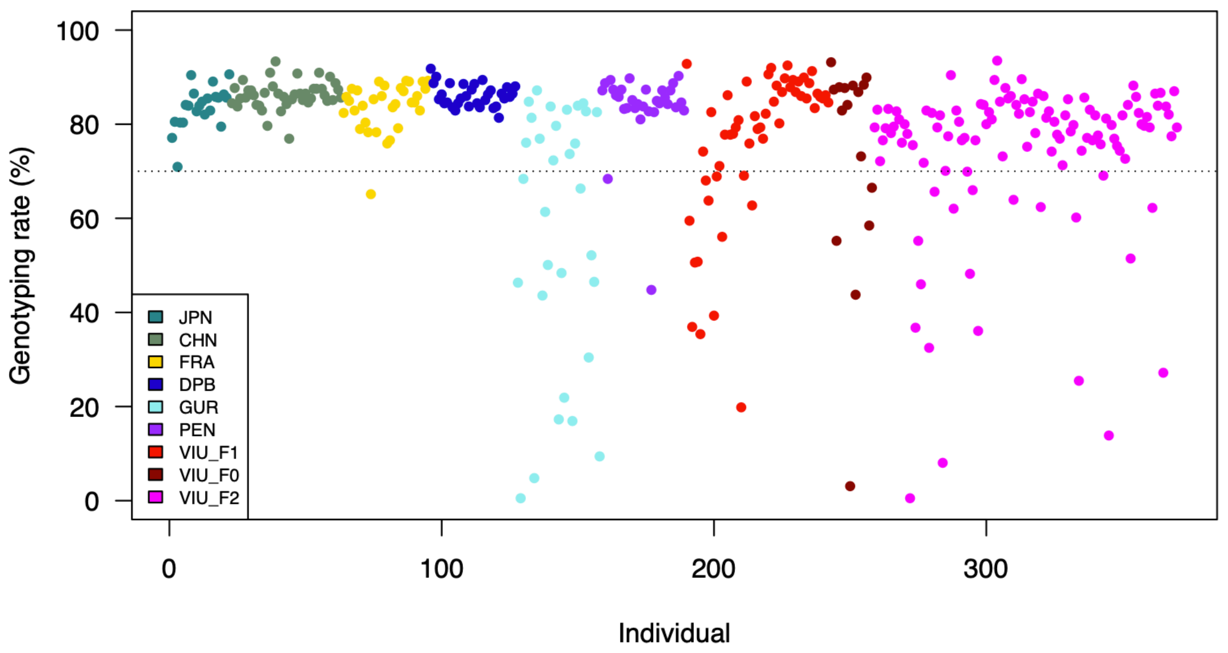


**Figure S3.** Per sample genotyping rate prior to any quality filters. Samples with more than 30% missing data (i.e., less than 70% genotyping rate) were removed from the analysis. Acronyms: JPN = Japan; CHN = China; FRA = France; DPB = Deep Bay; GUR = Guernsey; PEN = Pendrell; VIU = Vancouver Island University.

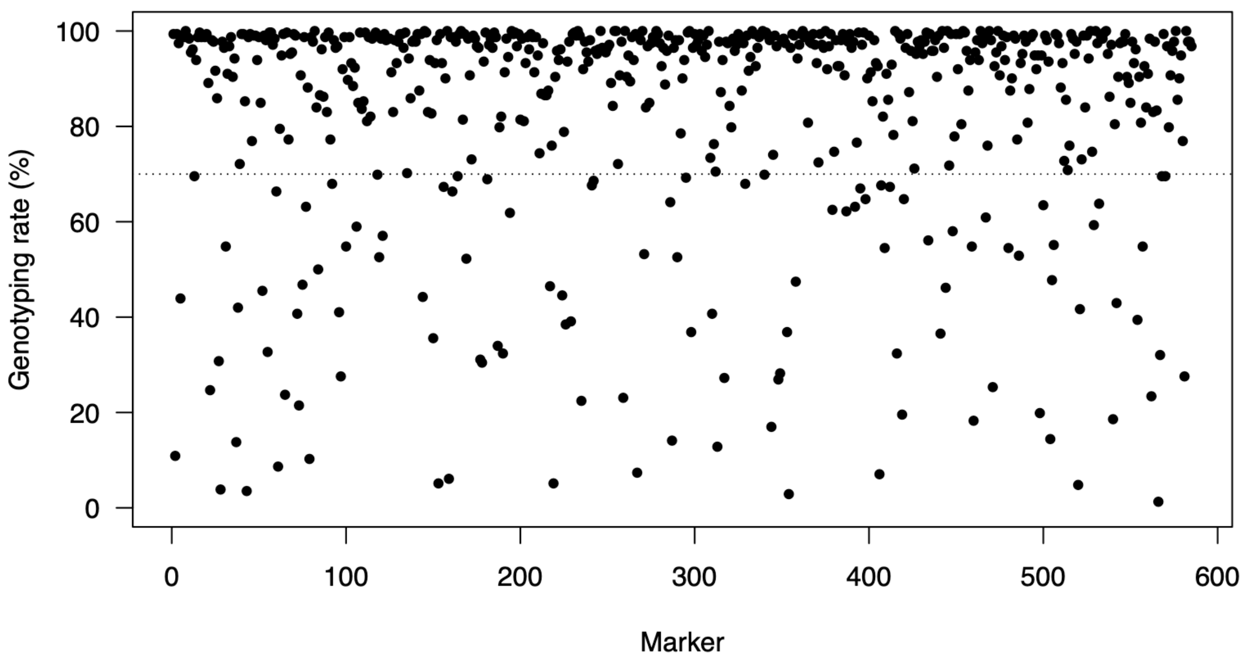


**Figure S4.** Genotyping rate per locus prior to quality filters. Loci with more than 30% missing data (i.e., less than 70% genotyping rate) were removed from the analysis.

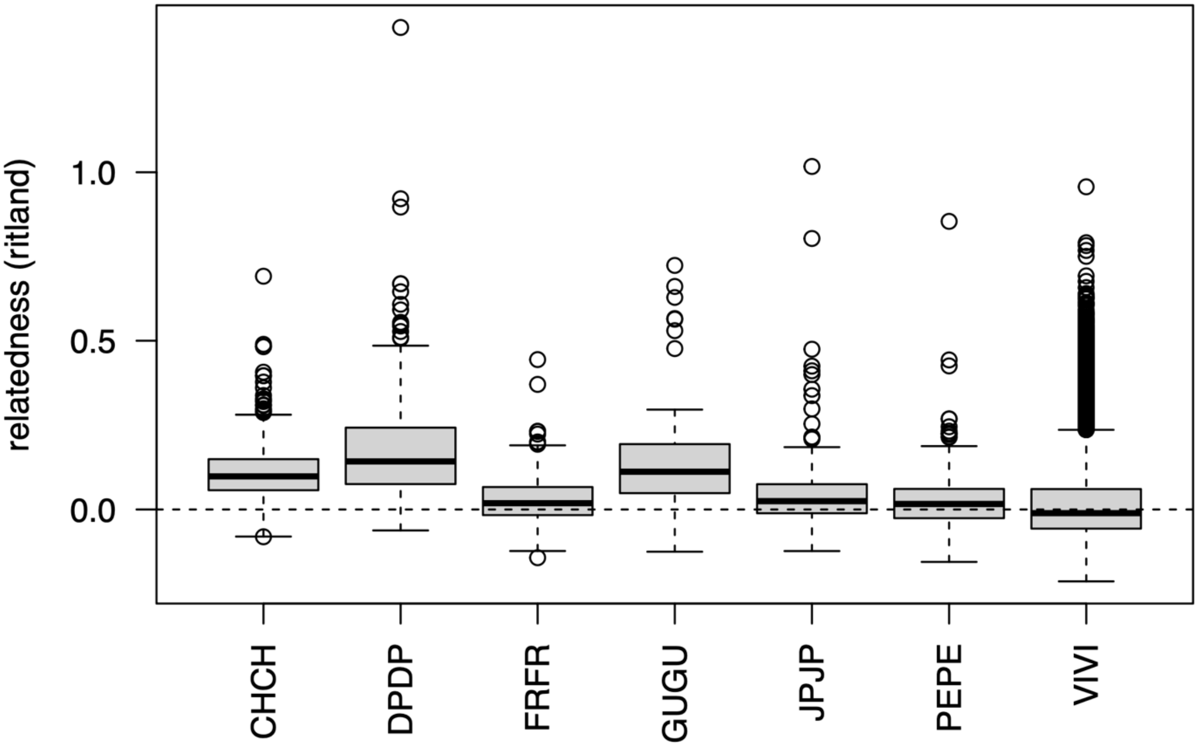


**Figure S5.** Inter-individual relatedness distributions between pairs of individuals within each population. Relatedness was estimated using the amplicon panel genotypes using the Ritland statistic (see Methods). Within population pair acronym: CHCH = China; DPDP = Deep Bay; FRFR = France; GUGU = Guernsey; JPJP = Japan; PEPE = Pendrell; and VIVI = VIU (all generations). Note: Guernsey, Deep Bay, and VIU samples are all hatchery or farm-based sampling events.

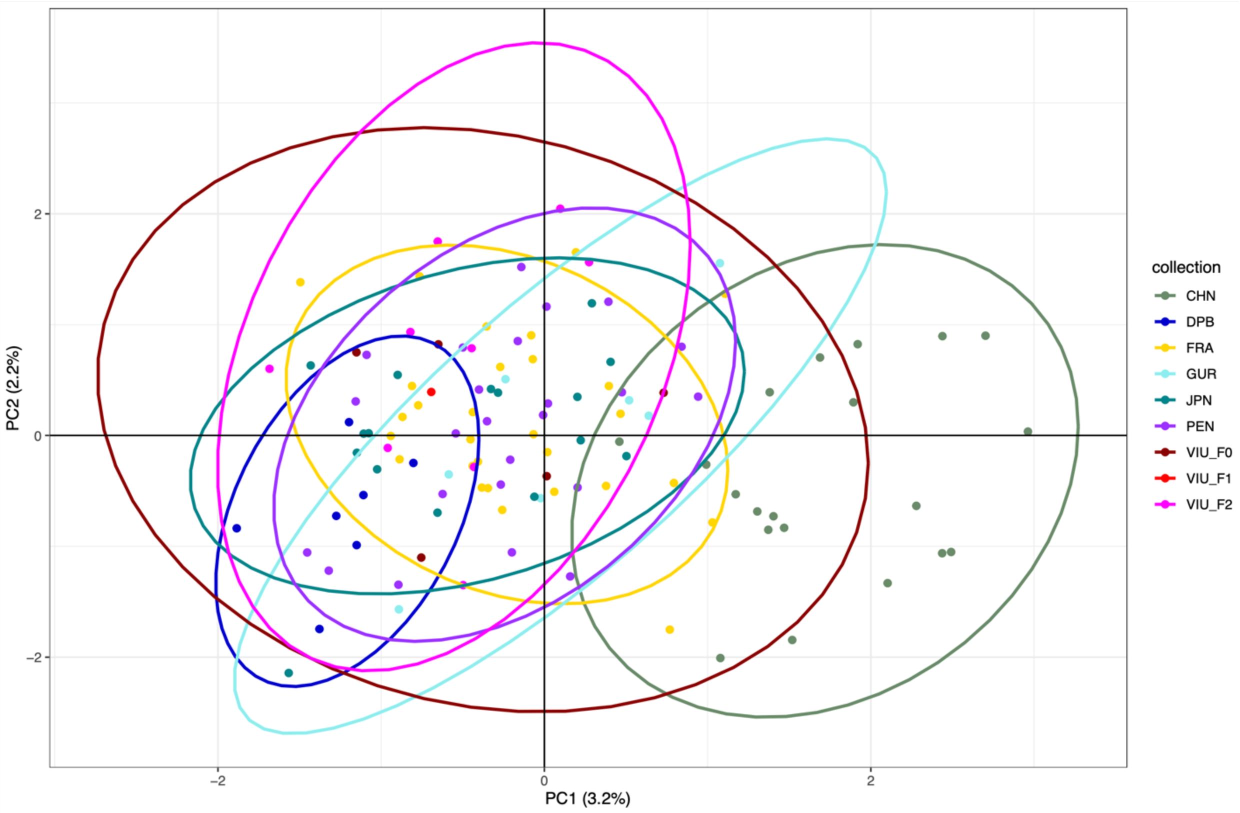


**Figure S6.** Principal components analysis (PCA) of pilot study samples clustered by filtered genotypes after estimated close relatives were removed (n = 118 individuals retained). Population abbreviations: VIU = Vancouver Island University (hatchery); DPB = Deep Bay (farm); PEN = Pendrell (naturalized); GUR = Guernsey (hatchery); FRA = France (naturalized); JPN = Japan (wild); CHN = China (wild).

**(A)**

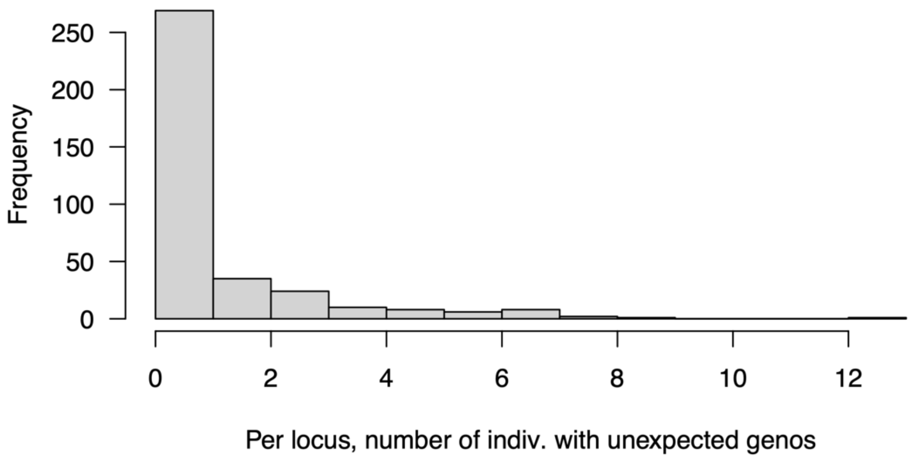


**(B)**

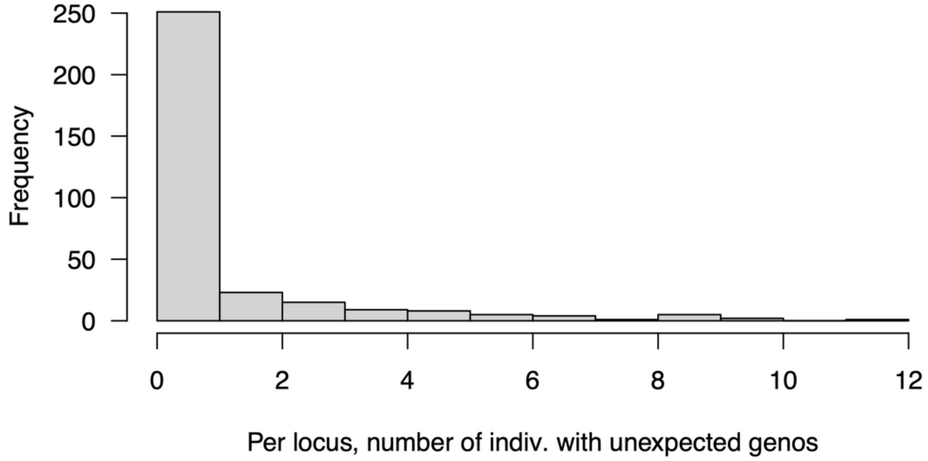


**Figure S7.** Per locus, number of offspring individuals that show unexpected genotypes based on empirically identified trios in the (A) pilot study VIU OFR5; or (B) MBP CHR8 families. Unexpected genotypes were identified by comparing per locus observed genotypes from offspring to the expected genotypes from the parents in the family.

**(A)**

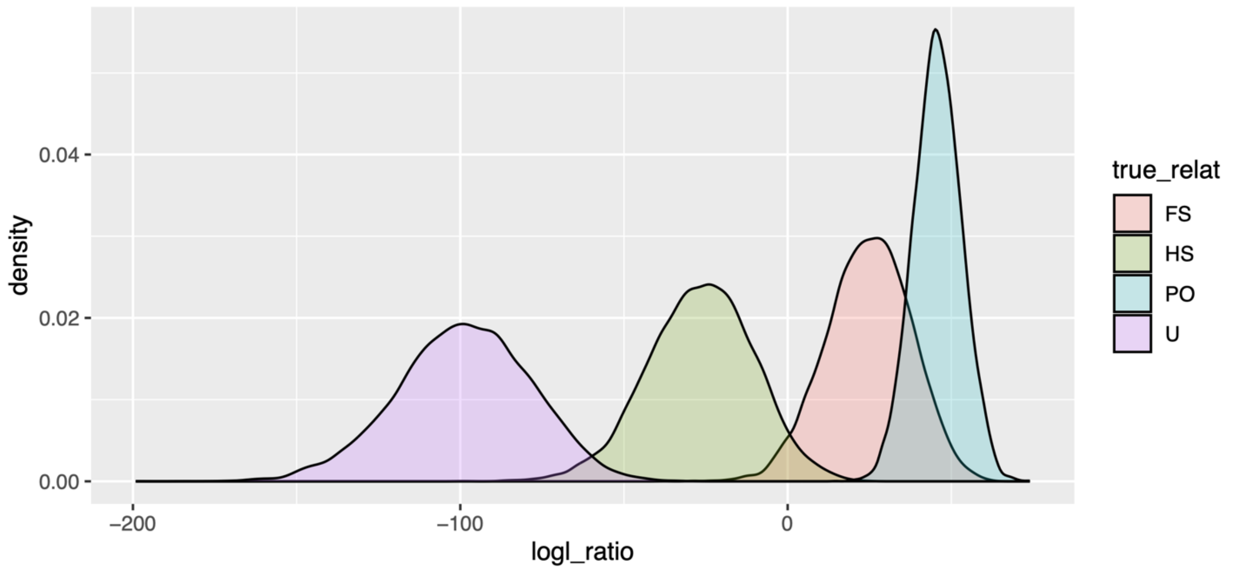


**(B)**

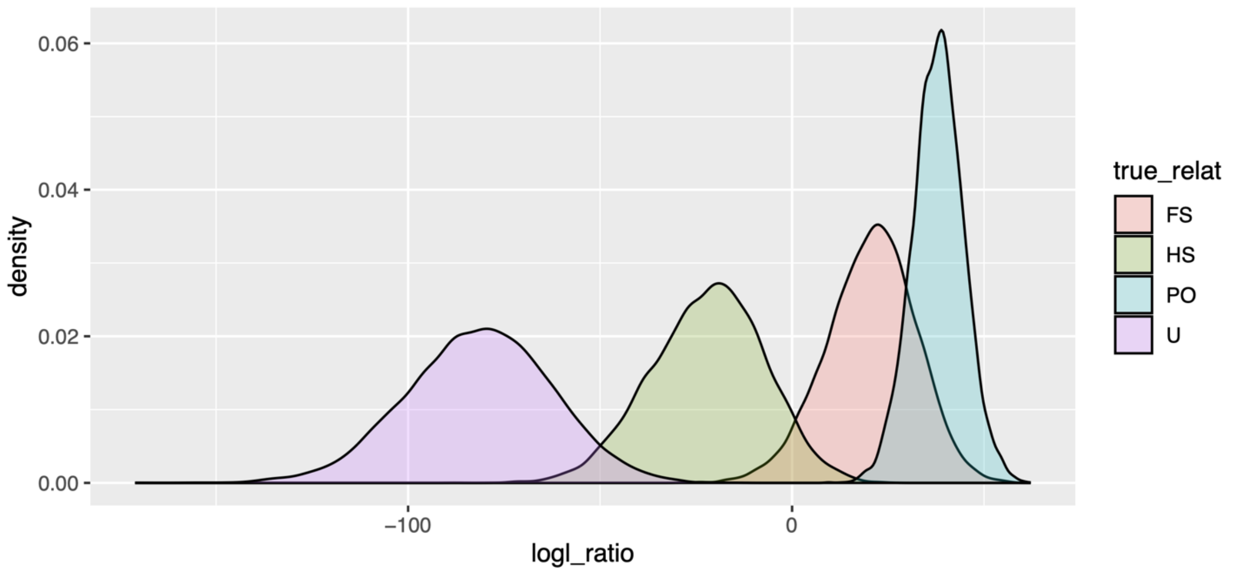


**Figure S8.** Log likelihood distributions for simulated relationships based on the amplicon panel genotypes after incompatible loci were removed. Relationships were simulated for parent-offspring (PO), full-sib (FS), half-sib (HS), and unknown (U) for datasets (A) VIU OFR5 cross (F1 vs. F2); and (B) MBP CHR8 families. Acronyms: VIU = Vancouver Island University; MBP CHR8 = Molluscan Broodstock Program chromosome 8; OFR = Oyster Family Run.

### Supplemental Tables

**Table S1.** Loci included in the Cgig_v.1.0 amplicon panel, including those submitted for design, those that were designed, those that passed quality control including minor allele frequency filters in the pilot dataset, and those that passed all filters, including removing incompatible genotype loci for both the VIU and the MBP CHR8 parentage datasets.

| Selection criterion | Markers (n) | Designed (n) | Pilot study pass initial filters | VIU parentage pass initial filters | MBP CHR8 study pass initial filters |
|---|---|---|---|---|---|
| $F_{ST}$ | 291 | 278 | 152 | 108 | 68 |
| *Heterozygosity* | 291 | 285 | 239 | 212 | 217 |
| $F_{ST}$ *and heterozygosity* | 9 | 9 | 7 | 3 | 4 |
| *Private alleles in DPB or GUR* | 20 | 20 | 11 | 5 | 0 |
| **Total** | **611** | **592** | **409** | **328** | **289** |

**Table S2.** Genetic differentiation ($F_{ST}$) between collections shown as lower (bottom left) and upper (top right) 95% confidence limits. $F_{ST}$ was evaluated from the amplicon panel data using 1000 bootstraps. PEN = Pendrell (naturalized); FRA = France (naturalized); JPN = Japan (wild); CHN = China (wild).

| | PEN | FRA | JPN | CHN |
|---|---|---|---|---|
| **PEN** | - | 0.0059 | 0.0117 | 0.0413 |
| **FRA** | -0.0014 | - | 0.0047 | 0.0361 |
| **JPN** | -0.0006 | -0.0044 | - | 0.0482 |
| **CHN** | 0.0211 | 0.0171 | 0.0231 | - |

**Table S3.** Private alleles identified empirically in the amplicon panel data when considering grouped collections from a similar area or translocation lineage (see *Methods*). Deep Bay (DPB) and Guernsey (GUR), both using different samples than those used for marker design identify high frequency private alleles that were included in the panel due to presence as private alleles in the marker discovery dataset (see *Results*).

| Grouping | Number of unique private alleles | Range of number of observations of the allele | Identities and counts of observations |
|---|---|---|---|
| JPN lineage | 7 | 1-5 | 576388.02 (5), 560208.02 (4), 576314.02 (3), 383999.02 (2), 417097.02 (1), 92747.02 (1), 713137.02 (1) |
| VIU | 3 | 1-8 | 590965.02 (8), 265979.02 (2), 93417.02 (1) |
| DPB | 3 | 2-8 | 39139.02* (8), 538043.02* (6), 8182.02 (2) |
| GUR | 2 | 3-9 | 336452.02* (9), 406807.02 (3) |
| CHN | 5 | 1-5 | 231185.02 (5), 442043.02 (2), 105579.02 (1), 344878.02 (1), 437400.02 (1) |

*****marker selected as private allele in different samples in RADseq dataset (Sutherland *et al.* 2020)
